## Supplementary material for "Abnormal AMPAR-mediated synaptic plasticity, cognitive and autistic-like behaviors in a missense *Fmr1* mutant mouse model of Fragile X syndrome"

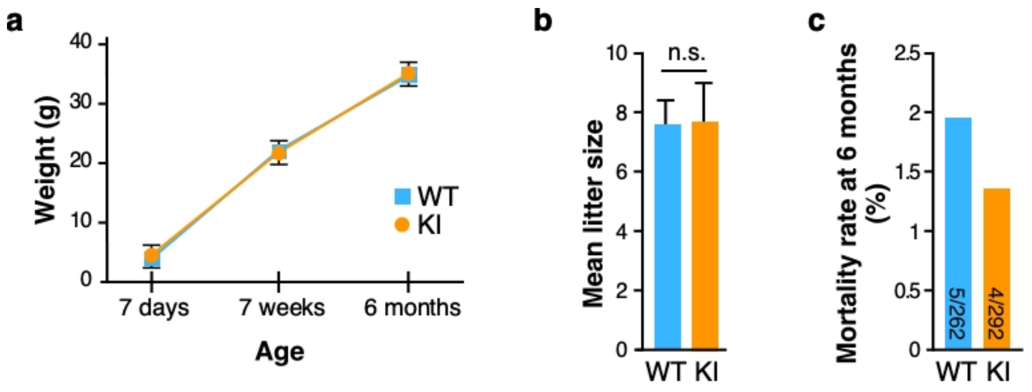


Supplementary figure 1

**Supplementary figure 1: *Fmr1$^{R138Q}$* mice does not show gross alterations.**

Quantification shows no significant differences between genotypes in the weight **(a)** of infant, adolescent and adult WT and *Fmr1$^{R138Q}$* KI mice, their fertility index **(b)** and mortality rate at 6 months **(c)**. Unpaired t-test. n.s., not significant.

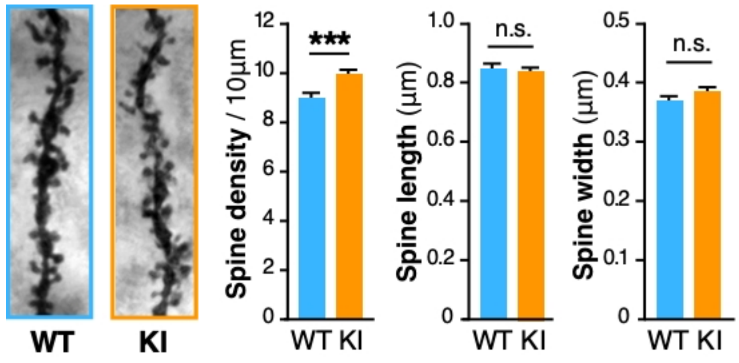


Supplementary figure 2

**Supplementary figure 2: The *Fmr1$^{R138Q}$* hippocampus exhibits increased dendritic spine density.**

Representative images of Golgi-stained apical secondary dendrites of CA1 hippocampal neurons from PND90 WT and *Fmr1$^{R138Q}$* KI littermates. Histograms show the density of spines, the spine length and width from WT and *Fmr1$^{R138Q}$* CA1 secondary dendrites. Bars represent mean ± s.e.m. N = 30 neurons per genotype from 3 independent experiments (1500-2000 spines analyzed per genotype). Mann-Whitney test; n.s., not significant. ***p<0.001.

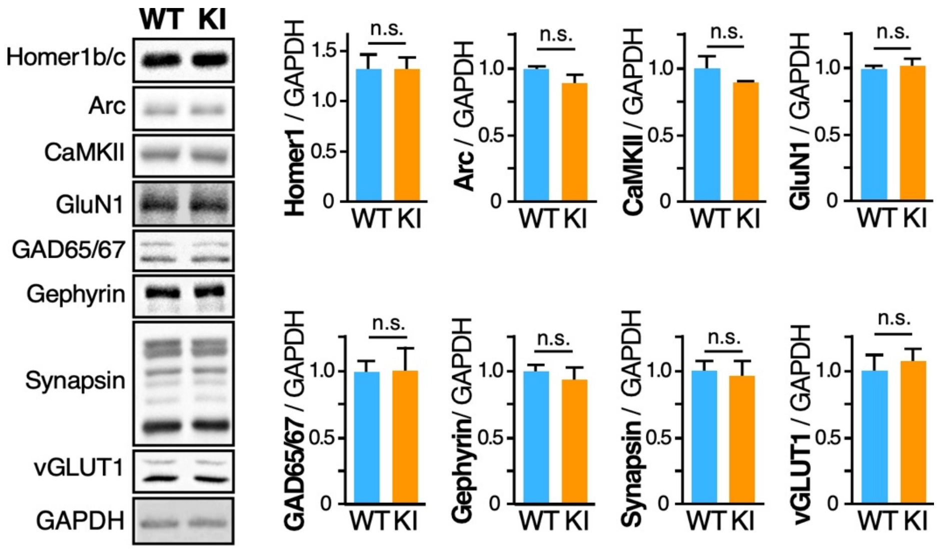


Supplementary figure 3

**Supplementary figure 3: Steady state levels of the indicated synaptic proteins in *Fmr1$^{R138Q}$* brain.** Representative immunoblots showing the levels of the indicated synaptic proteins in brain homogenates from WT and *Fmr1$^{R138Q}$* KI mice. GAPDH was used as a loading control as in figure 3a. Quantification shows the mean ± s.e.m. of the total levels of the indicated proteins. Unpaired t-test. n.s., not significant.

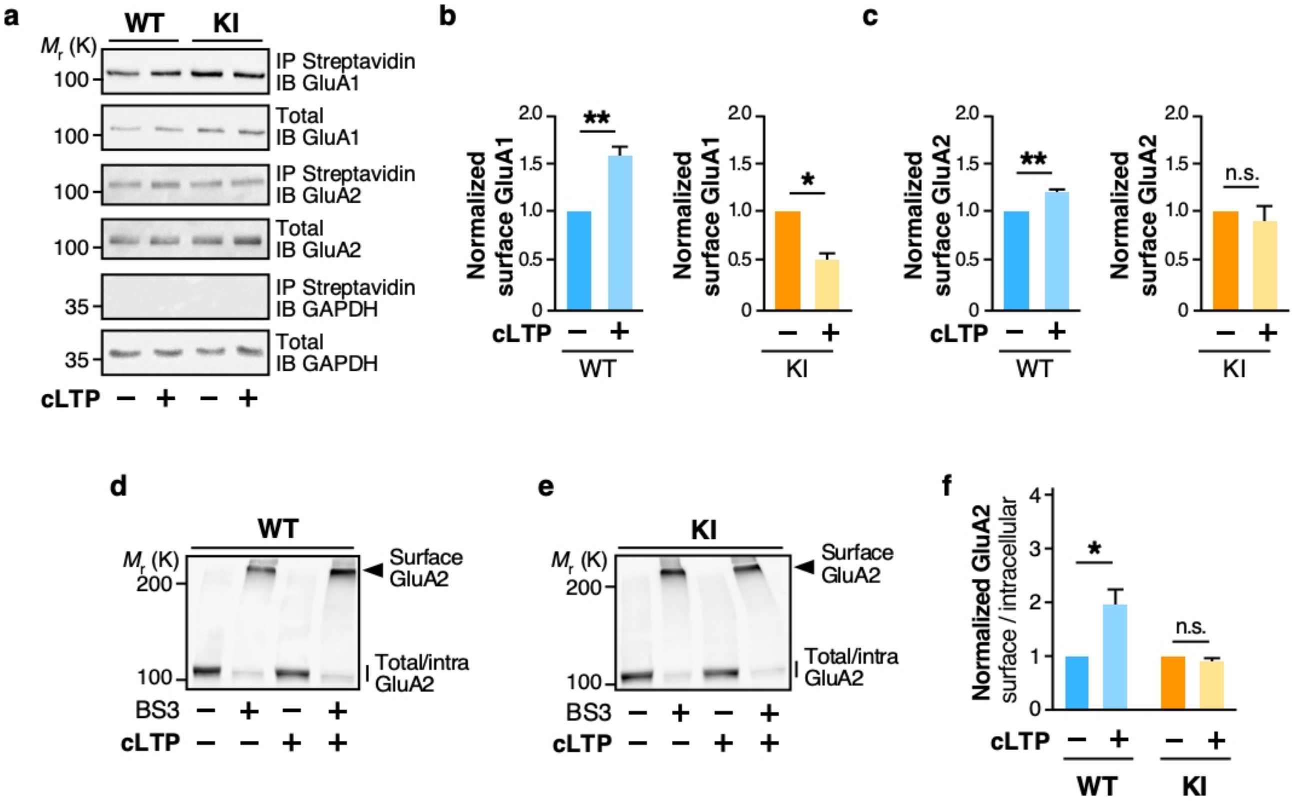


Supplementary figure 4

**Supplementary figure 4: Surface GluA2 levels are not increased in cLTP-induced *Fmr1*$^{R138Q}$ hippocampal cultures and slices.**

**a.** Representative immunoblots showing the surface expression of GluA1 and GluA2 in basal conditions and upon cLTP induction using biotinylation assays in cultured TTX-treated WT and *Fmr1*$^{R138Q}$ hippocampal neurons. Histograms show the mean ± s.e.m. of the surface levels of GluA1 **(b)** and GluA2 **(c)** from experiments in **(a)**. N=4 (GluA1) and 3 (GluA2) independent experiments. Ratio t-test. n.s., not significant. *p<0.05, **p<0.01. **d,e.** Representative immunoblots showing the surface expression of GluA2 in basal and cLTP-induced conditions in TTX-treated WT **(d)** and *Fmr1*$^{R138Q}$ **(e)** hippocampal slices using BS3 crosslinking assays. **f.** The surface/intracellular ratio in the WT was set to 1 and *Fmr1*$^{R138Q}$ KI values were calculated respective to the WT. Bars show the mean ± s.e.m. N = 5 independent experiments. Ratio t-test. *p=0.039. n.s., not significant.

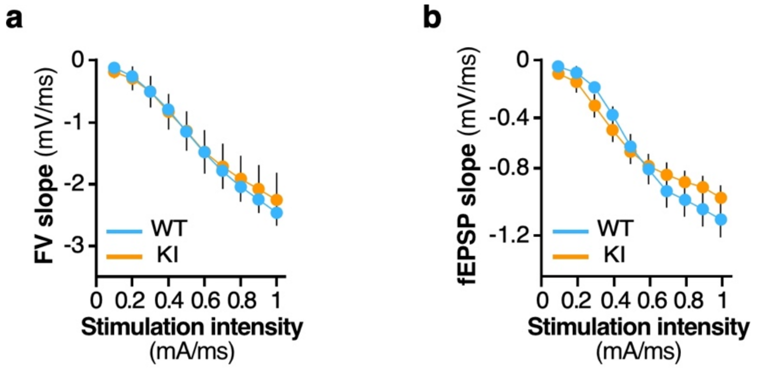


Supplementary figure 5

**Supplementary figure 5: *Fmr1$^{R138Q}$* mice show normal CA3 to CA1 synaptic connectivity.**

**a-b.** Input/output curves of Fiber Volley (FV, **a**) and postsynaptic responses (fEPSP, **b**) following Schaffer collaterals stimulations in WT and *Fmr1$^{R138Q}$* hippocampal slices.

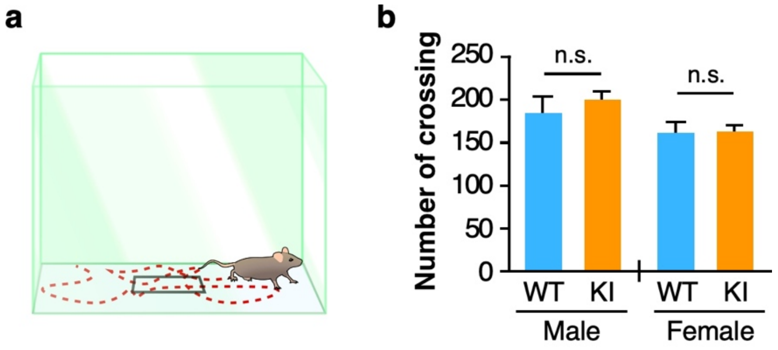


Supplementary figure 6

**Supplementary figure 6: *Fmr1$^{R138Q}$* mice do not show impaired locomotion.**

**a.** Scheme of the open field test used to assess locomotion. **b.** Quantification shows no significant differences between genotypes and genders. WT, N = 10 males, 9 females; KI *Fmr1$^{R138Q}$*, N = 10 males, 10 females. Two-way ANOVA with genotype and sex as factors followed by Newman-Keuls post-hoc test for individual group comparisons. n.s., not significant.

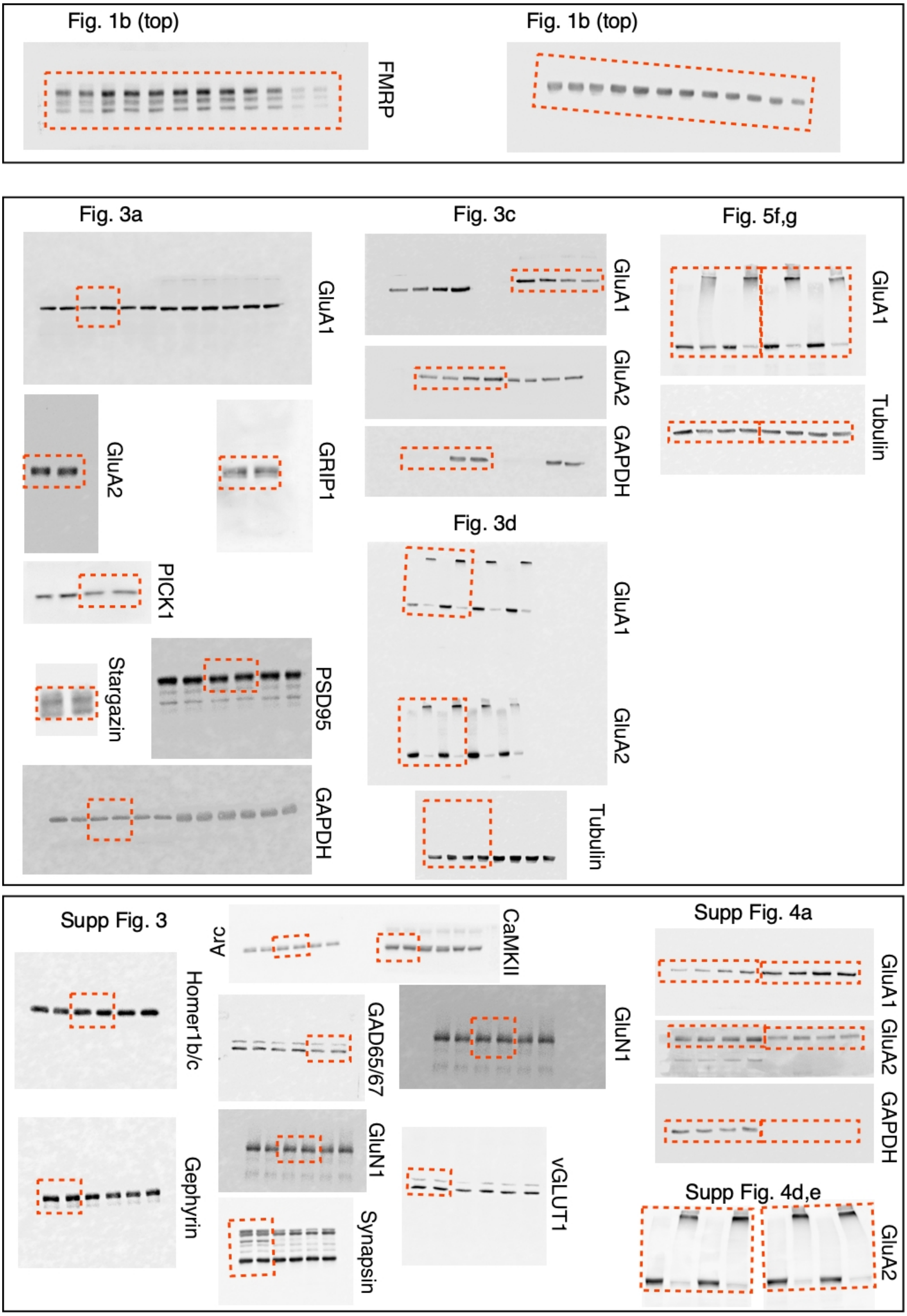


Supplementary figure 7

**Supplementary figure 7: Original uncropped blots.** Orange boxed regions represent the portion used in **figures 1, 3, 5** and **supplementary figures 3 and 4.**
